# A portable system for metagenomic analyses using nanopore-based sequencer and laptop computers can realize rapid on-site determination of bacterial compositions

**DOI:** 10.1101/101865

**Authors:** Satomi Mitsuhashi, Kirill Kryukov, So Nakagawa, Junko S Takeuchi, Yoshiki Shiraishi, Koichiro Asano, Tadashi Imanishi

**Affiliations:** Biomedical Informatics Laboratory, Department of Molecular Life Science, Tokai University School of Medicine, Isehara, Kanagawa 259-1193, Japan; Division of Pulmonary Medicine, Department of Medicine, Tokai University School of Medicine, Isehara, Kanagawa 259-1193, Japan; These authors contributed equally

## Abstract

We developed a portable system for metagenomic analyses consisting of nanopore technology-based sequencer, MinION, and laptop computers, and assessed its potential ability to determine bacterial compositions rapidly. We tested our protocols using mock bacterial community that contained equimolar 16S rDNA and a pleural effusion from a patient with empyema for time effectiveness and accuracy. MinION sequencing targeting 16S rDNA detected all of 20 bacteria present in the mock bacterial community. Time course analysis indicated that sequence data obtained during the first 5-minute sequencing were enough to detect all 20 bacteria species in the mock sample and determine their compositions with sufficient accuracy. Additionally, using a clinical sample extracted from the pleural effusion of a patient with empyema, we could identify major bacteria in a pleural effusion by rapid sequencing and analysis. All of these results are comparable to or even better than the conventional 16S rDNA sequencing results using IonPGM sequencer. Our results suggest that rapid sequencing and bacterial composition determination is possible within 2 hours.Our integrative system is applicable to rapid diagnostic tests for infectious diseases in near future.

## Introduction

Time is of importance when identifying pathogens in acute infectious diseases. In some critical conditions, such as bacteremia, starting antibiotic administration within 1 hour (h) is highly recommended (1). However, rapid pathogen identification is usually difficult as conventional bacterial culture takes more than a few days and some bacteria or fungi are difficult to grow in culture. Therefore, the initial choice of antibiotics is usually empirical. Establishing systems for rapid microorganism identification via metagenomic sequencing seems pertinent, especially for clinical use, to facilitate appropriate antibiotic treatment.

Most of the currently available sequencing technologies are basically not designed for rapid sequence analyses (2). These are based on several parallel fluorescence/proton scanning runs for obtaining large amounts of nucleotide sequence data, and it takes days to weeks to complete these runs However, the Oxford Nanopore Technology’s portable USB sequencer, MinION, produces nucleotide sequence data sequentially, enabling real-time metagenomic analysis. Moreover, MinION has many advantages such as simple sample preparation, portability, quick turnaround, maneuverability, and relative inexpensiveness. MinION has been in use for nearly 3 years since its release in 2014. There was a major update in its chemistry from R7 to R9 in May 2016, and the accuracy of sequencing has greatly improved. Rapid library preparation was also released in May 2016, which enables a 10-minute (min) library preparation from DNA to bacterial identification. This protocol only reads the template strand of the double-stranded DNA (“1D sequencing”) and was considered to be of a relatively low quality. In contrast, the original library preparation protocol produces both strands data (“2D sequencing”); 2D sequencing needs 90 min to prepare the library. R9 chemistry improved the usability of the data obtained from rapid 1D sequencing. These sequencing technology is increasingly being used for detecting pathogens in bacterial infection (3).

Data analysis for bacterial identification is also time-consuming, and may not be feasible in hospitals or diagnostic laboratories. *Centrifuge* is a novel microbial classification software that enables rapid and accurate identification of species (4) and can be run even on laptop computers. However, it is not clear what combination of these rapid sequencing provides a sufficiently quick and reliable bacterial detection system.

Here, we show that the combination of rapid sequencing using transposase-mediated library preparation for nearly full-length 16S rDNA amplicons, rapid 1D sequencing using R9 chemistry, 5-min sequencing data acquisition, local base-calling, and rapid analysis using *Centrifuge* on a laptop computer, enabled us to determine major bacteria within 2 h even in a small laboratory environment.

## Results

### Time-course analysis of MinION Rapid 1D sequence

We sequenced a 20 bacteria mock community (Supplemental Table 1) using various protocols to test their accuracies for detecting a variety of bacterial species. This mock bacterial community contains bacterial genomic DNA at an equimolar ratio for 16S rDNA for each species (5), and therefore, we expected to obtain the same number of reads for each bacteria species. Rapid 1D sequencing using SQK-RAD001 is designed for fast library preparation and utilizes transposase to fragment DNA, simultaneously attaching sequencing adapters that are necessary for sequencing to its free end. This is unsuitable for short amplicons; therefore, we amplified the almost full-length 16S rDNA (1399 bp) for MinION Rapid 1D sequencing (Figure 1B, Supplemental Tables 2-3) (8) We also tested the efficiency of our primer sets for 11 different bacterial species or strains that we could obtain from BEI resources (Supplemental Table 7). All 11 bacteria were also confirmed as amplified by our primer sets (Supplemental Figure 3A-B). On the other hand, MinION 2D sequencing and IonPGM sequencing were applied for the targeted sequences of 16S rDNA variable regions (Figure 1B, Supplemental Table 2).

**Figure 1.**
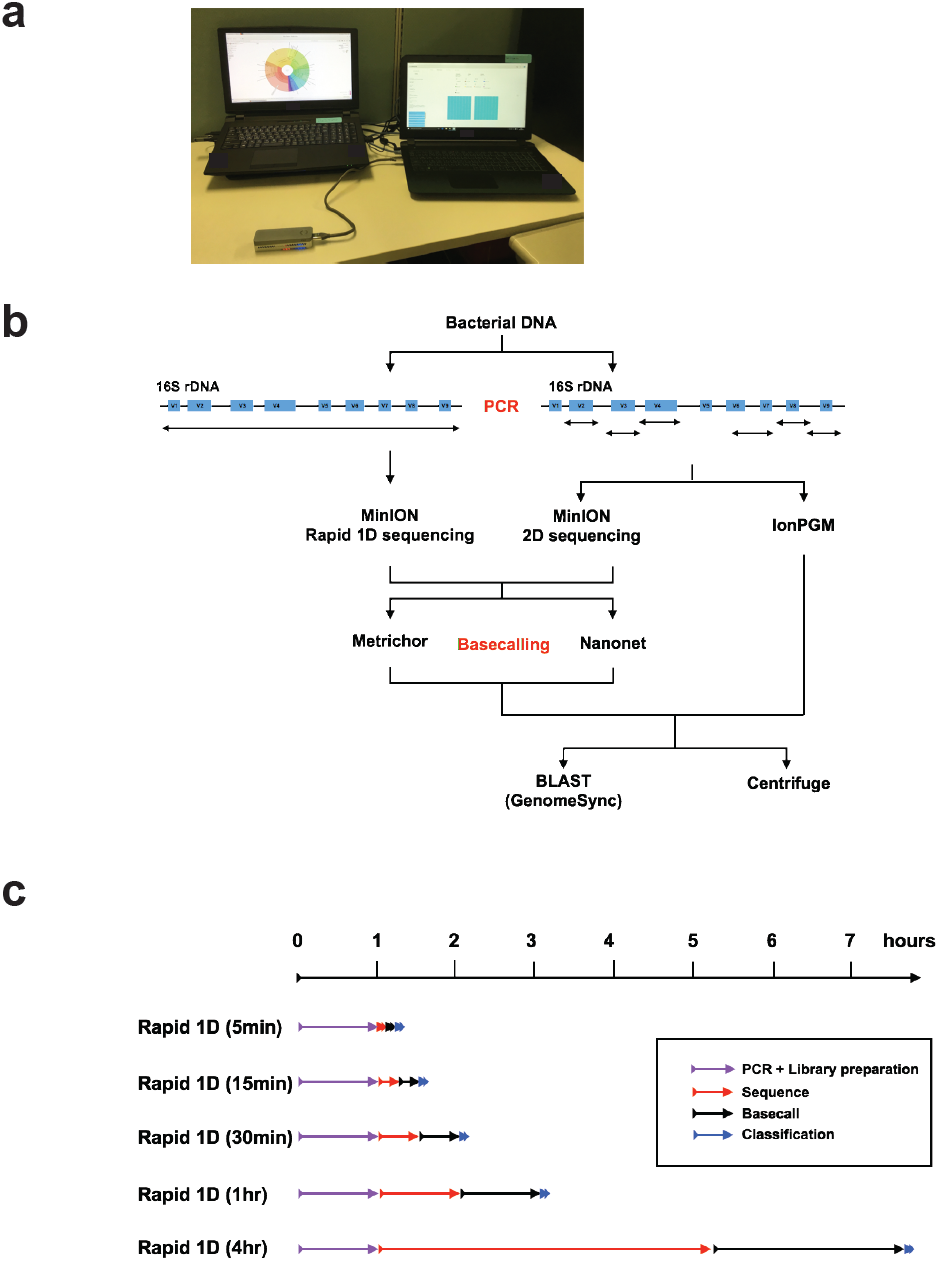
Portable system for bacterial determination and overview of our study (A) Picture of our system for bacterial composition determination. (B) Schematic outline of our experiment protocols. We used primer sets that amplified the same region of V2, V3, V4, V6-7, V8, and V9 for MinION 2D and IonPGM sequencing. For Rapid 1D sequencing, we amplified the almost full-length of the16S rRNA genes. For base-calling, an online software Metrichor and the local base-caller Nanonet were used. These results were analyzed both with *Centrifuge* and BLAST-based search method. (C) Rapid 1D sequencing data at different time points, 5 min, 15 min, 30 min, 1 h, and 4 h were collected and analyzed to assess time-effectiveness.

To assess time-effectiveness, sequencing data at five different time points (5 min, 15 min, 30 min, 1 h, and 4 h) from the beginning of MinION sequencing were utilized (Figure 1C) to compare different sequencing methods (Figure 1B). The difference between MinION 1D and 2D results may be primarily explained by the difference between 1D (only template) and 2D (both double-strands) sequencing, and/or by the difference of amplicon lengths (Figure 1B, Supplemental table 4). On the other hand, the difference between MinION 2D and IonPGM was mostly due to difference in sequencing methods because their target regions were identical to each other.

First, for the MinION sequencing data, the local base-calling Nanonet software was applied. Thereafter, the nucleotide sequences were searched with BLAST using our in-house genome database, GenomeSync. Overall, it was found that all sequencing protocols could identify all the 20 bacterial species in the reference mock community at both species and genus level (Figure 2A-F, Figure 3A-F). MinION 1D sequencing data showed the best sensitivity to 20 bacteria and less deviation from the expected read number both at the species and genus levels rather than IonPGM or 2D sequencing data (Figure 2A-F, Figure 3A-F, Supplemental table 5). Interestingly, 5-min sequencing data of 1,379 reads showed comparable results to those of 4 h sequencing of 24,202 reads. (Figures 2H and 3H, Table 1). Rapid 1D sequencing for 5 min showed 91% and 97% sensitivity at the species and genus levels, respectively (Figures 2H and 3H, Supplemental table 5).

**Figure 2.**
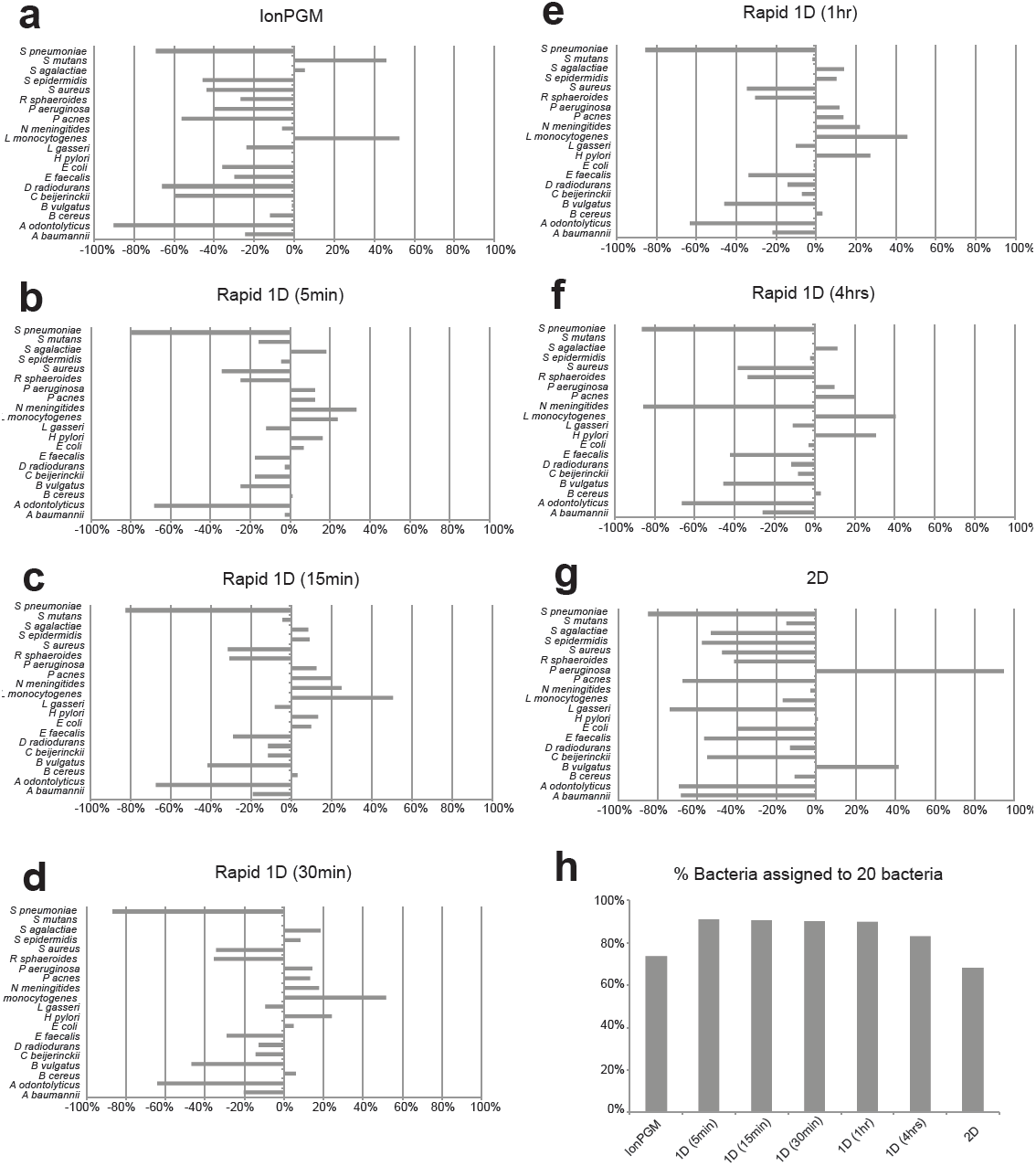
The bacterial classification for species level using the Nanonet base-caller and BLAST-based searches. Deviation from % expected reads to all the reads assigned to bacteria are shown (A-G). IonPGM sequencing for 16S rDNA (A). MinION Rapid 1D Sequencing of the almost full-length 16S rDNA amplicon for 5 min (B), 15 min (C), 30 min (D), 1 h (E), 4 h (F). MinION 2D sequencing for 16S rDNA (G). The percentage of reads assigned to any of the 20 bacteria among all reads classified as bacterial reads (H).

**Figure 3.**
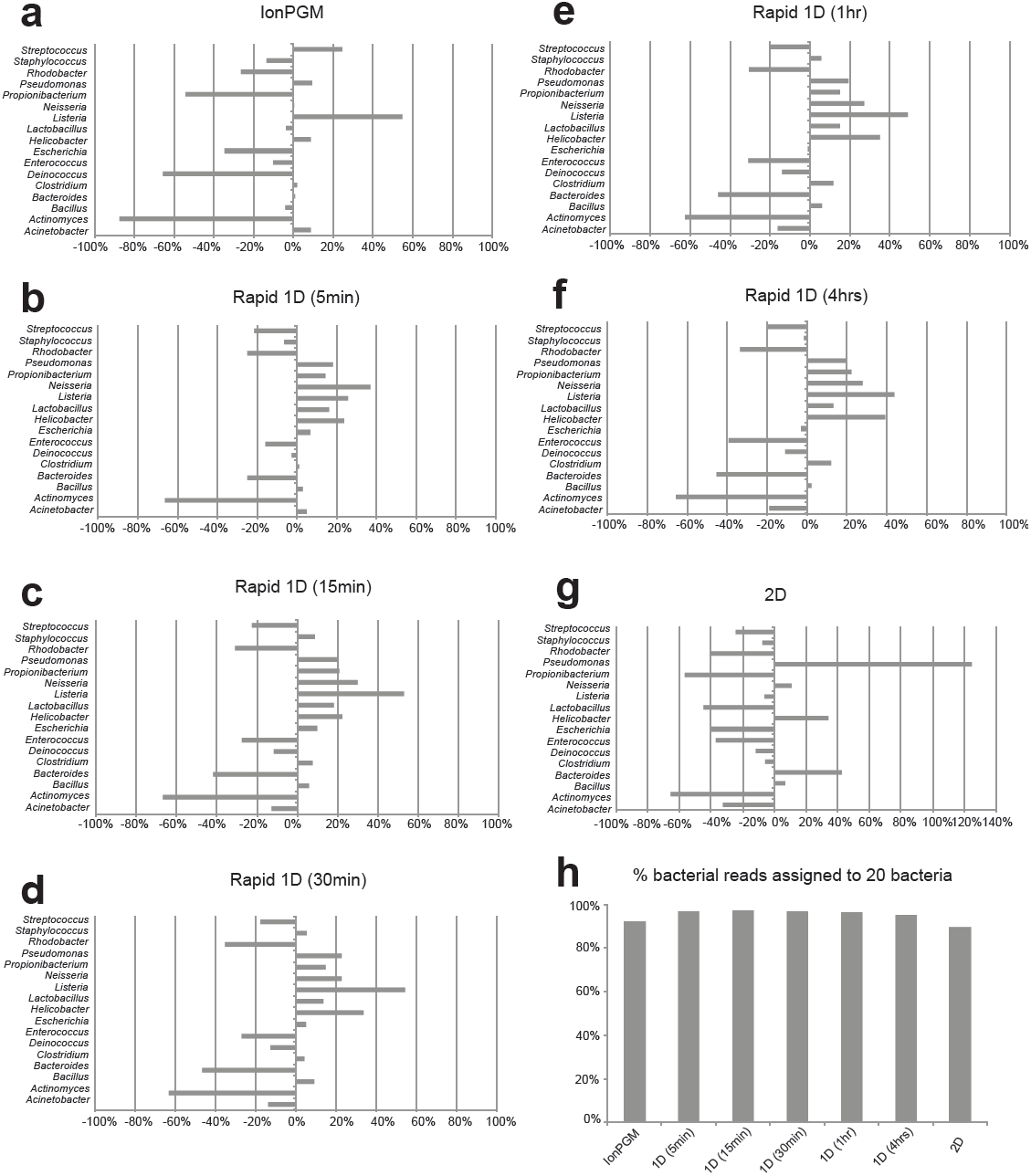
The bacterial classification at the genus level using the Nanonet base-caller and BLAST-based searches. Deviation from % expected reads to all the reads assigned to bacteria are shown (A-G). IonPGM sequencing for 16S rDNA (A). MinION Rapid 1D Sequencing of the almost full-length 16S rDNA amplicon for 5 min (B), 15 min (C), 30 min (D), 1 h (E), 4 h (F). MinION 2D sequencing for 16S rDNA (G). The percentage of reads assigned to any of the 20 bacteria among all reads classified as bacterial reads (H).

**Figure 4.**
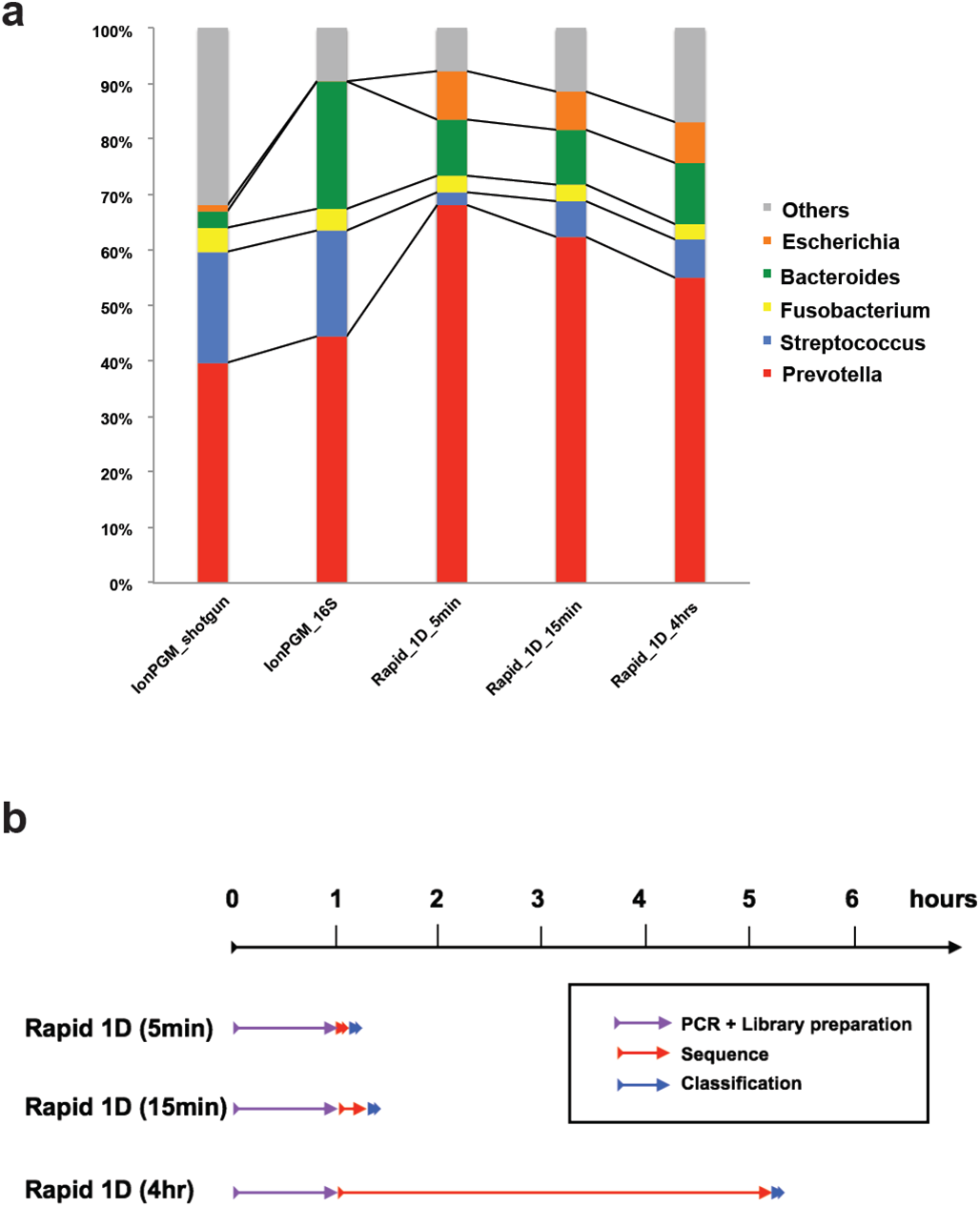
Clinical application of our system for determining bacterial composition in Pleural effusion from patient with empyema. (A) A pleural effusion sample sequenced using three different methods indicated that *Prevotella* was the predominant bacteria (B) Timescale for the experiment. Flow cell preparation took ~20 min for quality check and priming, which can be done during PCR.

We also tested nucleotide sequences base-called by Metrichor which uses cloud computing services through the internet (Figure 1B). In the Metrichor base-called data, 5-min Rapid 1D sequencing results showed 86% and 94% sensitivity at the species and genus level, respectively (Supplemental Figures 2H, 3H, Supplemental Table 5). This means that ~1000 reads are sufficient to detect 20 bacteria present in an equimolar manner. Overall, Nanonet base-calling showed better sensitivity for detecting 20 bacteria with less deviation compared to Metrichor (Supplemental Table 5). Furthermore, Nanonet is nearly 5 times faster than Metrichor (Table 1), although it requires a relatively high CPU power.

### Centrifuge analysis

BLAST-based search is time-consuming even when using computer clusters. To reduce the computational time for species detection, we tested a newly developed species classification suite, *Centrifuge* (4). At the species level, *Centrifuge* could identify almost all bacteria included in the mock community; however, *Actinomyces odontolyticus* was not identified by *Centrifuge*. On the other hand, at the genus level, all 20 bacteria were detected with a sensitivity of 63%, 68%, and 86% in the sequencing results of MinION Rapid 1D (5 min) and 2D, and IonPGM, respectively. *Actinomyces* was detected in 23%, 29%, and 15% of the expected *Actinomyces* reads in Rapid 1D sequencing (5 min), 2D sequencing, and IonPGM, respectively. BLAST analysis detected *Actinomyces* in 34%, 35%, and 12% of the reads in Rapid 1D sequencing (5 min), 2D sequencing, and IonPGM, respectively, suggesting that low detection of *Actinomyces* was not due to the *Centrifuge* software. Overall, *Centrifuge* exhibited less sensitivity and larger deviation compared to BLAST-based analysis, but the time spent on the bacterial classification was usually less than one min.

### Rapid 1D sequencing and analysis for Pleural effusion derived DNA

Finally, to examine whether this rapid sequencing is applicable for clinical samples, we sequenced total DNA samples extracted from the pleural effusion of a patient with empyema, in which microbiological examination identified *Streptococcus anginosus* group and unculturable Gram-negative rods. We compared the sequencing result between IonPGM sequencing and MinION rapid 1D sequencing. We also sequenced the same sample using IonPGM both with and without 16S rDNA amplification (16S rDNA sequencing and shotgun sequencing). As pleural effusion samples are rich in host-derived DNA, IonPGM shotgun sequencing showed only 0.37% of the total reads were classified as bacteria. All sequencing methods showed *Prevotella* as the major species in the pleural effusion in this patient (Figure 2A). *Prevotella* was detected in 40%, 44%, and 68% of all the bacterial reads in IonPGM shotgun sequencing, IonPGM 16S rDNA amplicon sequencing, and MinION rapid 1D sequencing (5 min), respectively, suggesting that *Prevotella* was the major species in the sample. Streptococcus was also detected by all the sequencing methods. *Centrifuge* provided the fastest classification with <20 sec for MinION data and <1 min for IonPGM data (Table 2). Total time from sample preparation to identification of bacteria was within 2 h (Figure 2B).

Further detailed analysis using BLAST with the GenomeSync database indicated that *Prevotella oris* was the major species in the sample. A BLAST search showed the composition of *P. oris* to total bacterial reads were 78%, 70%, and 85%, in IonPGM shotgun sequencing, IonPGM 16S rDNA amplicon sequencing, and MinION rapid 1D sequencing (5 min), respectively. However, *Centrifuge* could not detect *P. oris*, but instead classified the reads as other *Prevotella* species, indicating that this species genome may be not included in the *Centrifuge* genome index.

## Discussion

For clinical samples containing pathogenic bacteria, it would be informative to perform rapid and on-site sequencing to determine the first antibiotics of choice. Although this study is still in trial stages, our current report provides further evidence that a mock bacterial community was detected within 2 h using MinION sequencing and a data analysis platform on laptop computers. We performed data analysis on two different methods of bacterial classification: BLAST-based search and *Centrifuge*. *Centrifuge* is very rapid and could detect all the bacteria at the genus level; however, as our study suggests, it may misclassify some bacteria at the species level. One possible reason is that the prepared bacteria genome database for the *Centrifuge* program is relatively small (containing 4,078 bacterial genomes). It would be a powerful tool if this search method is combined with comprehensive genome data sets, or optimized for certain disease conditions. For example, a reference genome database for some common infectious disease such as pneumonia or bacterial meningitis may help rapid and accurate pathogen search. With such a system, it may be possible, in many cases, to promptly know what bacteria species is predominant in the specimen from infection focus. However, this approach may miss some rare pathogens. If this is the case, a second more intensive search based on a larger dataset may help increase the pathogen detection rate.

Although the time calculated here for computational analysis may differ from laboratory to laboratory because of computer performance, these data provide some insight for constructing a bacterial identification system for researchers. Our sequencing data of 20 reference mock bacterial community are available from DDBJ DRA database (DRA005399), and may be useful for researchers to examine analyses such as bacterial classification in their own environments.

Our result showed that rapid 1D sequencing using amplicons that covers almost full-length 16S rDNA has better sensitivity compared to sequencing protocols using short and multiple amplicons in BLAST analysis. This suggests that long read sequence is suitable for identification of the species. As MinION sequencer is capable of reading more than several thousands bases, it may be intriguing to target regions such as 23S rDNA which is longer than 16S rDNA or whole rDNA operon, which may show better resolution for bacterial identification.

We showed a rapid and accurate method for bacterial identification based on 16S rDNA PCR amplification; however, 16S rDNA sequencing has some limitations (i.e., drug resistance bacteria identification). Schmidt et al. reported that they could detect bacteria drug-resistant genes by directly sequencing DNA from a heavily infected urine samples using MinION (3). They used bacteria-rich urine samples, therefore, human DNA contamination was minimized. This method may be difficult to apply for bacteria-scarce samples, such as blood specimens from infected patients, as some reports showed that it may need >30 million reads (12). However, currently, new sequencing technologies are continuously being improved regarding data size and accuracy. It may be possible to sequence bacteria-scarce clinical samples without PCR amplification when MinION capacity improves in the near future.

## Conclusions

Our results suggest that a 2-hour rapid determination of the bacterial composition using the MinION sequencer and laptop computers is feasible and applicable for clinical use as a diagnostic support tool in hospitals or small laboratories. Further improvements regarding the computational bacterial identification method may provide a better resolution at the species level.

## Materials and Methods

### Bacterial DNA

Genomic DNA from 20 different bacteria was obtained from Biodefense and Emerging Infectious Research (BEI) Resources (http://www.beiresources.org) of the American Type Culture Collection (ATCC) (Manassas, VA, USA) (5). Mock Microbial Community B (BEI catalog number HM-782D) contains 20 different bacterial strains with an equal molar of 16S rDNA copy for each organism. Double strand DNA concentration was measured by Qbit Fluorimeter using dsDNA HS Assay kit (Thermo Fisher Scientific, MA, USA).

### 16S rDNA amplicons and PCR

The 16S primers used in this study are shown in Supplemental Table 2. Six sets of primers for short amplicon sequencing were designed for 16S rDNA variable regions (V2, V3, V4, V6-7, V8, and V9), for 2D sequencing. The pooled PCR product was subjected to 2D DNA library preparation. Primers S-D-Bact-0008-c-S-20 and S-D-Bact-1391-a-A-17, which almost cover the full length of 16S rDNA were used for 1D sequencing (6–9). One nanogram of DNA was used for PCR. KAPA HiFi HotStart ReadyMix (KAPA Biosystems, MA, USA) and Agencourt AMPure XP system (Beckman Coulter Genomics, CA, USA) were used for PCR and PCR product purification, respectively. The PCR condition is shown in Supplemental Table 3.

### IonPGM sequencing for 16S and data analysis

Bacterial 16S rDNA regions were amplified by PCR using 2 primer pools covering variable regions (V2, V3, V4, V6-7, V8, and V9) from the Ion 16S™ Metagenomics Kit. Thereafter, emulsion PCR was performed using the Ion OneTouch™ System that was loaded onto an Ion 316™ Chip. It was sequenced using an Ion PGM™ with HiQ chemistry according to the manufacturer’s protocol (Thermo Fisher Scientific, MA, USA). Fastq files were generated by Ion Reporter System (Thermo Fisher Scientific, MA, USA), which were subjected to data analysis.

### Amplicon DNA library preparation and DNA sequencing using MinION

Library preparation was performed using SQK-NSK007 Nano Sequencing Kit R9 version (Oxford Nanopore Technologies, Oxford, UK) and Rapid Sequencing Kit SQK-RAD001 (Oxford Nanopore Technologies, Oxford, UK) using 1 µg and 200 ng amplicon DNA, for 2D amplicon sequencing and rapid 1D sequencing, respectively, according to the manufacturer’s protocol. MinION sequencing was performed using MinION Mk1 b sequencer and FLO-MIN104 flow cells. Raw data (fast5 files) were obtained using MinKNOW software ver. 1.0.2 (Oxford Nanopore Technologies, Oxford, UK).

### Data Analysis

Base-calling was performed using Nanonet and Metrichor softwares, both developed by Oxford Nanopore Technologies. Nanonet is a local (i.e., no internet-connection required) base-caller based on recurrent neural network (RNN) algorithms, while Metrichor is a cloud-based software that uploads row output files produced by MinION (fast5 files) to the server and downloads base-called fast5 files. We used Nanonet 1D sequence with default parameters. We also used Metrichor’s 1D Base-calling (FLO_MIN105 250bms) for 1D sequencing fast5 data and 2D Base-calling RNN (SQK-MAP007) for 2D sequencing fast5 data. For base-called Metrichor fast5 files, quality scoring and conversion to FASTA format were performed using poretools ver 0.6.0 (10).

### Time-course dependent bacterial identification using BLAST and *Centrifuge*

Rapid bacterial identification was performed with the *Centrifuge* software using bacterial, viral, and human genome dataset (4). Identification of bacteria was also performed using BLAST search with the GenomeSync database (http://genomesync.org). Representative bacterial and fungal genomes as well as the human genome were used. Taxa were determined using our in-house script, then visualized using Krona Chart (11). We performed bacterial composition analysis with the following dataset; MinION 1D sequence reads with Nanonet and Metrichor during the first 5 min, 15 min, 30 min, 1 h, and 4 h from the time sequencing was started. We also compared the 1D sequencing results with those from MinION 2D sequencing and IonPGM 16S Metagenomics sequencing.

### Laptop Computers

Two laptop computers were used for the analysis. One was for MinION sequencing and Metrichor base-calling (OS, Windows 10; CPU, Intel Core i7 6700HQ; memory, 8 GB; storage, 960 GB SSD); the other was for Nanonet base-calling, running poretools and bacterial identification by *Centrifuge* (OS, CentOS 7; CPU, Intel Core i7 6700K; memory, 32 GB; storage, 1 TB SSD) (Figure 1A).

### DNA preparation from the pleural effusion in a patient with empyema

A pleural effusion sample was collected from a 77-year-old, male patient with empyema for diagnostic purposes with written informed consent. This patient had a history of diabetes, cerebral infarction and gingivitis. The collected sample was centrifuged at 50 *x G* for 10 min to remove human white blood cells and was then re-centrifuged at 15,500 *x G* for 10 min to pellet the bacteria. After removing the supernatant, the cells were re-suspended with Lysis Buffer (20 mM TrisHCl, pH 8.0, 2 mM EDTA, 20 µg/µl Lysozyme, 1.2% TritonX) at 37°C for 30 min and subjected to DNA extraction using DNeasy mini kit (Qiagen) according to the protocol. Extracted DNA was used for IonPGM sequencing (Thermo Fisher Scientific, MA, USA) and MinION 1D sequencing using Rapid 1D sequencing kit with S-D-Bact-0008-c-S-20 and S-D-Bact-1391-a-A-17 primers.

### Pleural effusion sample sequencing and analysis

MinION 1D sequencing was performed using Rapid Sequencing Kit SQK-RAD001 (Oxford Nanopore Technologies, Oxford, UK) using 200 ng amplicon DNA. MinION Mk1 b sequencer and FLO-MIN105 flow cell were used for sequencing. In this sequencing run we used MinKNOW software ver. 1.1.17. This version allows local base-calling for 1D sequencing data using protocol script NC_48Hr_Sequencing_Run_FLO_MIN104_plus_1D_Basecalling to minimize the time for the analysis, although this may produce different base-calling results compared to Metrichor or Nanonet. Bacterial identification was performed using BLAST and *Centrifuge* as described above and were compared with two sequencers. IonPGM sequencing using the same DNA was performed both for 16S rDNA amplicons and Covaris-fragmented DNA (shotgun sequencing). For 16S rDNA amplicon sequencing, the above-mentioned 16S Metagenomics Kit was used (Thermo Fisher Scientific, MA, USA). Shotgun sequencing was performed according to the manufacture’s protocol (Thermo Fisher Scientific, MA, USA). Briefly, 100 ng of DNA was fragmented using Covaris sonicator (Covaris, MA, USA) and was then subjected to library preparation using a fragmentation kit (Thermo Fisher Scientific, MA, USA). Adapter-ligated DNA fragment was size-selected using eGel (Thermo Fisher Scientific, MA, USA).

### Ethics

This study was carried out with the approval of the Institutional Review Board for Clinical Research, Tokai University School of Medicine (14R220).

## Acknowledgments

The authors thank for the Support Center for Medical Research and Education, Tokai University, for providing technical support. We also thank Meiko Takeshita Miho Sera, Kentaro Mamiya and Takuya Habara for technical assistance. The following reagent was obtained through BEI Resources, NIAID, NIH as part of the Human Microbiome Project: Genomic DNA from Microbial Mock Community B (Even, Low Concentration), v5.1L, for 16S rRNA Gene Sequencing, HM-782D.

## Author Contributions

SM, KK, SN, JST, and TI designed the study. SM and JST collected experimental materials. YS and KA collected the clinical materials. SM conducted experiments. SM, KK and SN analyzed and interpreted the data. SM, KK, SN and KA wrote the manuscript. TI approved the manuscript to be submitted.

## Web Resources

The URLs for data presented are as follows: GenomeSync http://genomesync.org.

## Data access

The 20 bacterial mock community data was submitted to the DDBJ DRA database under accession number DRA005399.

